# Cold, but not for long enough: first insights into the tolerance to subzero temperatures of the invasive amphipod *Dikerogammarus villosus*

**DOI:** 10.1101/2023.10.30.562454

**Authors:** Serena Mucciolo, Krzysztof Podwysocki, Andrea Desiderato

## Abstract

*Dikerogammarus villosus* is an invasive Ponto-Caspian amphipod, known to have colonized most of the major European freshwater bodies in the last few decades. Several studies assessing its tolerance related to higher temperatures have been performed; however, no records are present about its response to subzero temperatures. Therefore, in this study, we wanted to experimentally assess for the first time the tolerance of *D. villosus* to temperatures below 0°C. The animals were exposed to 0°C, -8°C, and room temperature for 2 and 24 hours, tracking their possible movements during the ice melting. Only long exposures at -8°C appeared to be fatal. These results provide new insights for more accurate predictive models regarding the future expansion of this invader.

## Introduction

One of the main sources of European freshwater non-indigenous and invasive species is the Ponto-Caspian region, which is characterized by a very dynamic geological history (several sea-level changes) and consequently fluctuating environmental stressors, such as salinity, temperature, and oxygen^1^. Several Ponto-Caspian species were intentionally introduced into other regions of Europe for aquaculture, but were also unintentionally aided by anthropogenic modifications of the natural topography of rivers and channels, connecting regions otherwise isolated and eventually opening new pathways to their spread.

For instance, from this region is *Dikerogammarus villosus* (Sowinsky, 1894), a freshwater gammaridean amphipod, considered among the worst invasive species in Europe^2^. It was able to colonize most of the major European freshwater bodies in only a few decades, replacing many native amphipod species, including previous invaders, and preying on other taxa, such as egg fishes and tadpoles^3–5^. Its invasion success relies on a generally broader physiological niche (i.e., euryoecious), faster population growth (e.g., high fecundity, early maturity), a diverse diet (i.e., generalist), and aggressive competitive behaviour^6–10^.

Temperature is one of the major challenges affecting organisms from the cellular to the population level, influencing their biological activities (e.g., metabolism, development), population dynamics (e.g., reproduction), and eventually their distribution (e.g., ^11,12^). Animals usually display a diverse range of sensitivity to thermal fluctuations, as well as a combination of adaptations (e.g., specific proteins, thermoregulating behaviour) to buffer this stress. In the last few decades, climate change has led to an increase in studies assessing the sensitivity of several taxa to high temperatures, as well as its impact on native communities in relation to the spread of invasive species^13,14^. Melting ice sheets, the consequent variation in sea surface salinity and ocean circulation, seem to be linked to the climate regime shifts, which may lead to abrupt temperature changes with quicker and shorter warm temperature periods and longer cooling terms^15^.

The knowledge about amphipod response to lower temperatures is scattered. For instance, the survival of some circumpolar gammarid species at subzero temperatures and after being trapped inside the ice was well documented. Similarly, the effect of the ice contact after the exposure to low temperatures and cold acclimation (i.e., +3 and -2°C) was assessed in the subterranean *Niphargus rhenorhodanensis* Schellenberg, 1937, and compared to the surface-dwelling *Gammarus fossarum* Koch, 1836^16–18^. The metabolic rate was also assessed in *D. villosus*, which showed better performances (i.e., reduced energy expenditure) at lower temperatures (i.e., 5°C, 15°C) when compared with two European native gammarids (*G. fossarum*, and *G. roeselii* Gervais, 1835)^12^. Moreover, *D. villosus* seems to tolerate freezing, being able to survive at subzero temperatures for a few hours, trapped below or inside a layer of ice (Bącela-Spychalska and Mamos’ pers. comm.). Thus, the aim of this study was to experimentally investigate for the first time the sensitivity of *D. villosus* to subzero temperatures to provide novel information about its potential range expansion.

## Results and discussions

After two hours at –8°C, all the specimens were completely frozen. However, after thawing, their movements were comparable to those of the control and 0°C treatments (**Fig. 1**). It is worth stressing that the animals successfully underwent an abrupt change in temperature (∽ 30 °C of difference). This is the first record of an amphipod from a temperate region surviving complete freezing. Although the longer exposure resulted to be fatal (i.e., no movements recorded), the ability of *Dikerogammarus villosus* to survive at cytosol-freezing temperatures is relevant for its eco-physiological implications.

**Figure 1.**
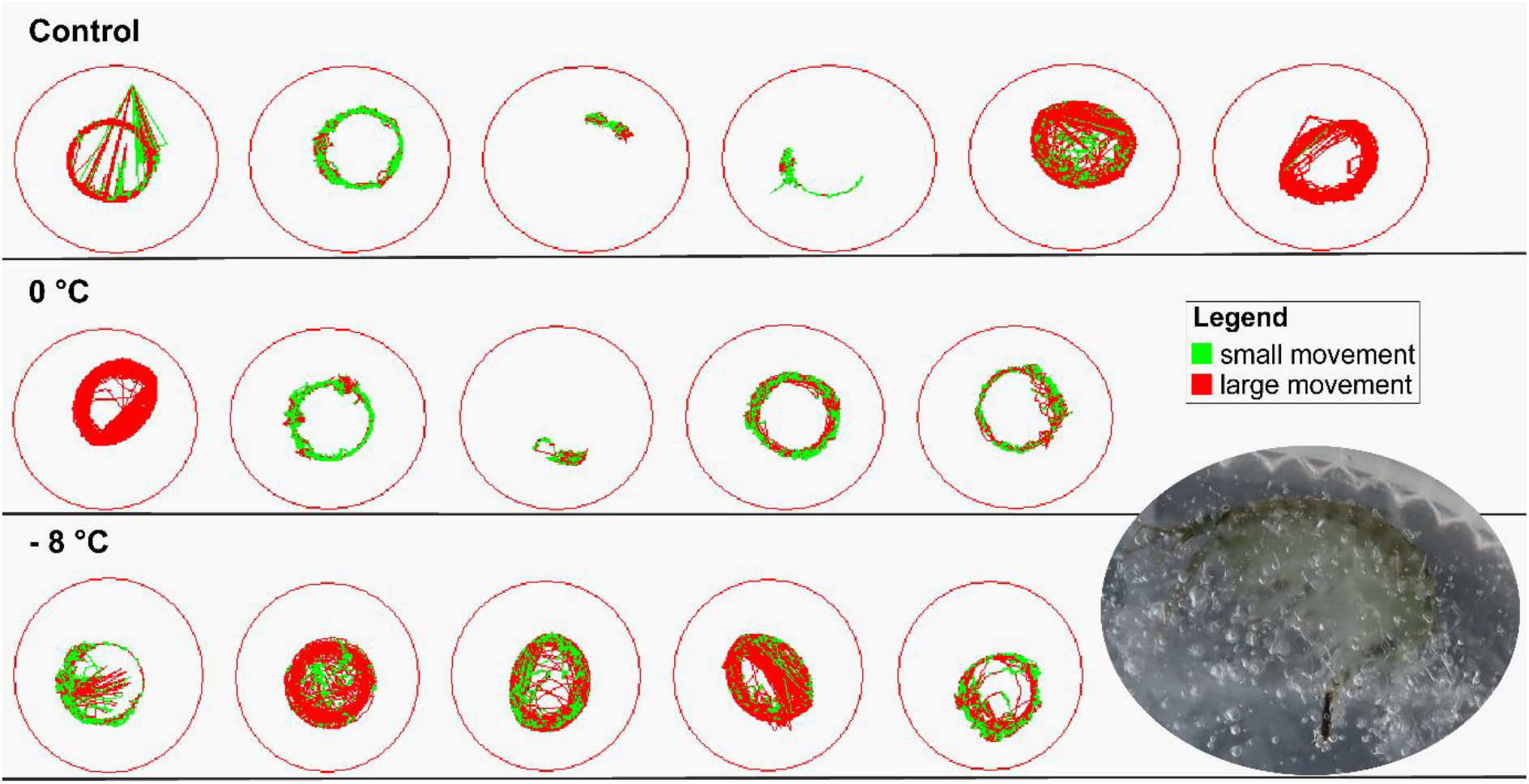
*Dikerogammarus villosus* activity recordings after 2 hours of exposure, in each experimental condition. In the photo, a frozen specimen after 2 hours of exposure at -8°C.

The repeated measures during the three hours after the treatments were never significant (**Table 1**), meaning that the ice thawed during the first 10 minutes and the amphipods started to move as soon as they were freed, as clearly visible in the video (**Suppl. video S1, Suppl. table S1**).

**Table 1.** Results of the analysis of deviance table of the GLMM. Significant differences are indicated in bold.

|  |  | Small distance |  |  | Large distance |  |  |
| --- | --- | --- | --- | --- | --- | --- | --- |
| Control treatment | Coefficient | Chisq | Df | Pr(>Chisq) | Chisq | Df | Pr(>Chisq) |
|  | Time | 2.32 | 2 | 0.31 | **99.56** | **2** | **<0.001** |
| Subzero treatment | Treatment | 4.70 | 2 | 0.10 | **19.40** | **2** | **<0.001** |
|  | Time | **6.14** | **1** | **0.01** | **17.19** | **1** | **<0.001** |
|  | Interval | 0.17 | 1 | 0.68 | 1.30 | 1 | 0.25 |
|  | Treatment:Time | **15.00** | **2** | **<0.001** | **97.00** | **2** | **<0.001** |
|  |  | Small duration |  |  | Large duration |  |  |
| Control treatment | Coefficient | Chisq | Df | Pr(>Chisq) | Chisq | Df | Pr(>Chisq) |
|  | Time | **6.94** | **2** | **0.03** | **60.03** | **2** | **<0.001** |
| Subzero treatment | Treatment | 4.10 | 2 | 0.13 | **17.27** | **2** | **<0.001** |
|  | Time | **5.25** | **1** | **0.02** | **11.88** | **1** | **<0.001** |
|  | Interval | 0.23 | 1 | 0.63 | 0.82 | 1 | 0.36 |
|  | Treatment:Time | **12.92** | **2** | **0.00** | **70.01** | **2** | **<0.001** |

**Table** 298 **Table 1.** Results of the analysis of deviance table of the GLMM. Significant differences are 299 indicated in bold.
|  |  | **Small distance** |  |  | **Large distance** |  |  |
| --- | --- | --- | --- | --- | --- | --- | --- |
| **Control treatment** | Coefficient | Chisq | Df | Pr(>Chisq) | Chisq | Df | Pr(>Chisq) |
|  | Time | 2.32 | 2 | 0.31 | **99.56** | **2** | **<0.001** |
| **Subzero treatment** | Treatment | 4.70 | 2 | 0.10 | **19.40** | **2** | **<0.001** |
|  | Time | **6.14** | **1** | **0.01** | **17.19** | **1** | **<0.001** |
|  | Interval | 0.17 | 1 | 0.68 | 1.30 | 1 | 0.25 |
|  | Treatment:Time | **15.00** | **2** | **<0.001** | **97.00** | **2** | **<0.001** |
|  |  | **Small duration** |  |  | **Large duration** |  |  |
| **Control treatment** | Coefficient | Chisq | Df | Pr(>Chisq) | Chisq | Df | Pr(>Chisq) |
|  | Time | **6.94** | **2** | **0.03** | **60.03** | **2** | **<0.001** |
| **Subzero treatment** | Treatment | 4.10 | 2 | 0.13 | **17.27** | **2** | **<0.001** |
|  | Time | **5.25** | **1** | **0.02** | **11.88** | **1** | **<0.001** |
|  | Interval | 0.23 | 1 | 0.63 | 0.82 | 1 | 0.36 |
|  | Treatment:Time | **12.92** | **2** | **0.00** | **70.01** | **2** | **<0.001** |

The specimens in the control treatment displayed a significant decrease in their large movements after the first two hours (both distances and durations), but not in the small ones (**Table 1**). This can be attributed to a general high energetic consumption at 23 °C coupled with the relative stress related to the continuous movement and the lack of a shelter^19^. Moreover, the general preference of *D. villosus* for lower temperatures (i.e., 5 – 10°C) - reflected also in its spatial distribution in Europe, and a limited tolerance to higher temperatures (∽ 20-25°C) has already been reported in several studies^12,20,21^. Our results suggest the potential further spread of this invasive species to higher latitudes, boosted also by the recent finding of this species in Sweden^22^.

The individual effect for the small movements was particularly high with estimated standard deviations exceeding the fixed terms for both variables (i.e., Marginal R^2^ / Conditional R^2^: 0.466 / 0.933 - distances, 0.244 / 0.934 - durations; table SX). On the other end, this effect was smaller for larger movements (i.e., Marginal R2 / Conditional R2: 0.868 / 0.922 - distances), 0.642 / 0.878; **Suppl. table S2**).

The tolerance to very low temperature is the result of a combination of physiological and autecological adaptations. Concerning the physiological adaptations, previous studies have already pointed out in this species the presence of proteins potentially involved in thermal stress (e.g., ^23^) as well as the presence of a high glycogen reserve^21^, which is very common in several taxa - both vertebrates and invertebrates - adapted to extreme freezing temperatures (e.g., ^16,24,25^). Several polar and subpolar amphipod species adopt a behavioural thermoregulation, escaping the ice by migrating into deeper waters^18^; similarly, *D. villosus* frequently occurs in deeper water during winter. Active ice avoidance is indeed a common strategy because once freezing begins, ice crystals, extracellular or intracellular, create and propagate quickly^26^. Indeed, the temperature of -8°C seemed to be critical also for the circumpolar *G. oceanicus*, when completely frozen without allowing the animals to move in the water column^18^. Thermal tolerance is also often associated with salinity tolerance, and therefore, several euryhaline organisms are often as well eurythermal (e.g., ^27,28^). For instance, polar gammarids can tolerate higher salinities when exposed to very low temperatures, switching from osmoregulation to osmoconformity^18^. As *Dikerogammarus villosus* is an euryhaline species, being able to tolerate salinities < 25 PSU^28^ its osmotic response may vary with temperature changes. This study represents the first attempt to experimentally determine the physiological thermal niche of *D. villosus* for subzero temperatures, showing how this species bears an even larger unexpected set of traits that may contribute to explain and possibly help hinder its expansion.

## Methods

### Sampling, experimental set-up and statistical analysis

To avoid potential population-driven biases in the response, specimens of *Dikerogammarus villosus* were collected from the Oder River, in two locations: Zdzieszowice (50.4117°N, 18.1071°E) and Brzeg (50.8605°N, 17.4665°E). In both sites the water temperature was 24 ± 0.5 °C and the salinity was 1 ± 0.2 PSU. The animals were transported in 3 l buckets with ∽200 ml of river water and *Salix alba* leaves conditioned for more than 10 weeks. The buckets were cooled in styro-boxes with ice to avoid high temperatures during transport. In the laboratory, animals were acclimatized in 60 x 40 x 10 cm trays filled with 7 l of aerated water, with 20 washed stones of an average diameter of 5 cm to serve as shelters for the animals, and constant aeration and a 12:12 h light:dark period. Animals were fed daily with frozen chironomid larvae. Only specimens in good condition after the acclimatization (i.e., moving without evident damage or missing appendages) and non-ovigerous females were used. Three temperatures were chosen for the experiment: 0°C, to simulate the water temperatures reached during the winter in Poland^29^; -8°C, considering the subzero temperature at which the cytosol starts freezing, i.e., ∽ -5°C^30^; room temperature of 23 ± 1 °C as a control. To avoid a non-controllable time effect, all the treatments were carried out simultaneously, although in different environments or equipment (fridge Vestfrost model FKG371, portable freezer VigoCool V42W). Two exposure times, 2 h and 24 h, were chosen to assess the effect in both the short and longer term. Each cold treatment had five replicates per exposure time, while six were used for the control (i.e., 16 specimens per time). To measure the amphipods’ activity, a ZebraCube ZEB437, complemented with ZebraLab v3.12 software (Viewpoint Life Technology, Lyon, France), was used. The type of activity was classified as small and large with default settings measuring the amount of distance (mm) and their duration (s). The movements were measured cumulatively every 600 s for up to 3 hours (i.e., 16 times). During the first two hours of the experiment, the movements of the specimens of the control treatment were measured as an estimation of reference activity, complementing the 12 replicates with an additional four specimens (i.e., 16 specimens measured for two hours – 12 times). The videos were further checked manually to confirm the actual movement of the specimens.

### Statistical analyses

Extreme outliers were a priori removed (i.e., large distances > 10,500 mm, small distances > 6,100 mm, large durations > 500 s). First, a model comparing the control (room temperature) treatments along the whole experiment (i.e., first two hours, after them, and after 24 hours) was fitted for each of the four dependent variables (i.e., small and large distances or durations). Given the non-normal nature of the data with an exponential distribution and a high proportion of zeros, a Tweedie generalized linear mixed model was used with the glmmTMB function of the homonymous package^31^. As the specimens were repeatedly measured, they were included in the model as a random effect (i.e., 1|amphipodID). The effect of the different temperatures was modeled using a similar approach but including the interaction with the treatments and the different measurement times (i.e., sets of 600 s) as a covariate. The Wald chi-square test was computed for each model through analysis of deviance with the ANOVA function in the “car” package^32^. Pairwise comparison was performed with the emmeans function and Bonferroni adjustments via the homonymous package^33^. All analyses were performed in R software 4.3.0^34^.

## Acknowledgements

The authors would like to thank Tomasz Mamos and Karolina Bącela-Spychalska for their personal observations about the gammarid response to subzero temperatures prior to the experiments, łukasz Pułaski for the fruitful conversation and the suggestion to carry out the experiment, Marco Vito Guglielmi and Jakub Bienias for the help during the sampling. The sampling was carried out under the IDUB project (#B2311001000195.07) from the University of Lodz.

## Author contributions

SM: conceptualization, funding acquisition, investigation, data curation, visualization, writing – original draft, writing – review and editing. KP: investigation, data curation, software, writing – review and editing. AD: conceptualization, project administration, formal analysis, methodology, visualization, software, supervision, validation, writing – original draft, writing – review and editing. All authors have read and approved the final manuscript.

## Data availability statement

All data generated and analysed during this study are included in this published article and its Supplementary Material files.

